# PSAURON: a tool for assessing protein annotation across a broad range of species

**DOI:** 10.1101/2024.05.15.594385

**Authors:** Markus J. Sommer, Aleksey V. Zimin, Steven L. Salzberg

## Abstract

Evaluating the accuracy of protein-coding sequences in genome annotations is a challenging problem for which there is no broadly applicable solution. In this manuscript we introduce PSAURON (Protein Sequence Assessment Using a Reference ORF Network), a novel software tool developed to assess the quality of protein-coding gene annotations. Utilizing a machine learning model trained on a diverse dataset from over 1000 plant and animal genomes, PSAURON assigns a score to coding DNA or protein sequence that reflects the likelihood that the sequence is a genuine protein coding region. PSAURON scores can be used for genome-wide protein annotation assessment as well as the rapid identification of potentially spurious annotated proteins. Validation against established benchmarks demonstrates PSAURON’s effectiveness and correlation with recognized measures of protein quality, highlighting its potential use as a general-purpose method to evaluate gene annotation. PSAURON is open source and freely available at https://github.com/salzberg-lab/PSAURON.

**One-Sentence Summary:** PSAURON is a machine learning-based tool for rapid assessment of protein coding gene annotation.

## INTRODUCTION

Eukaryotic genomes continue to be sequenced, assembled, and annotated at a rapid pace ^1^. Accurate annotation of these genomes remains a significant challenge, with errors potentially resulting in profound negative implications for downstream analyses. The quality of genome annotation, particularly of protein-coding gene annotation, is crucial for understanding gene function and for a wide range of applications in biotechnology and medicine ^2^. While many metrics have been developed to assess the quality of genome assemblies ^3–5^, few methods currently exist to assess the accuracy of annotation.

Here we introduce PSAURON (pronounced like “Sauron” ^6^) a novel software tool that accurately scores predicted protein sequences and enables the genome-wide evaluation of protein-coding gene annotation. Using a temporal convolutional network (TCN), a machine-learning model that we previously applied to prokaryotic genome annotation ^7^, PSAURON analyzes coding sequences to assess the likelihood that a given annotation correctly identifies protein-coding genes. We trained the PSAURON TCN on a comprehensive dataset of protein sequences from a broad range of plant and animal genomes, enabling it to recognize general patterns indicative of protein coding sequences without the need for retraining on any individual species. One benefit of PSAURON lies in its ability to quickly assess whole-genome annotations with minimal computational requirements, providing researchers with a score that reflects the confidence in annotation accuracy for each protein coding sequence. This scoring system is designed to highlight potential inaccuracies in annotations, facilitating the identification of sequences that may require further investigation.

We validated PSAURON against well-established reference datasets and protein quality metrics to demonstrate its effectiveness in distinguishing between high- and low-quality gene annotations. As more genomes continue to be sequenced, the need for accurate and efficient tools for assessment of proteins will become increasingly critical. PSAURON represents a step forward in this regard, offering to the genomics community an efficient solution for quality control in protein-coding gene annotation.

## METHODS

### Model Architecture

The temporal convolutional network (TCN) architecture used here is based on one developed by Bai et al. ^8^ and modified to work with protein sequence data in Balrog, a bacterial gene finder ^7^. Figure 1 highlights some unique aspects of the TCN that make it amenable to working with amino acid sequence data, particularly its ability to produce a single likelihood score for any length protein.

**Figure 1:**
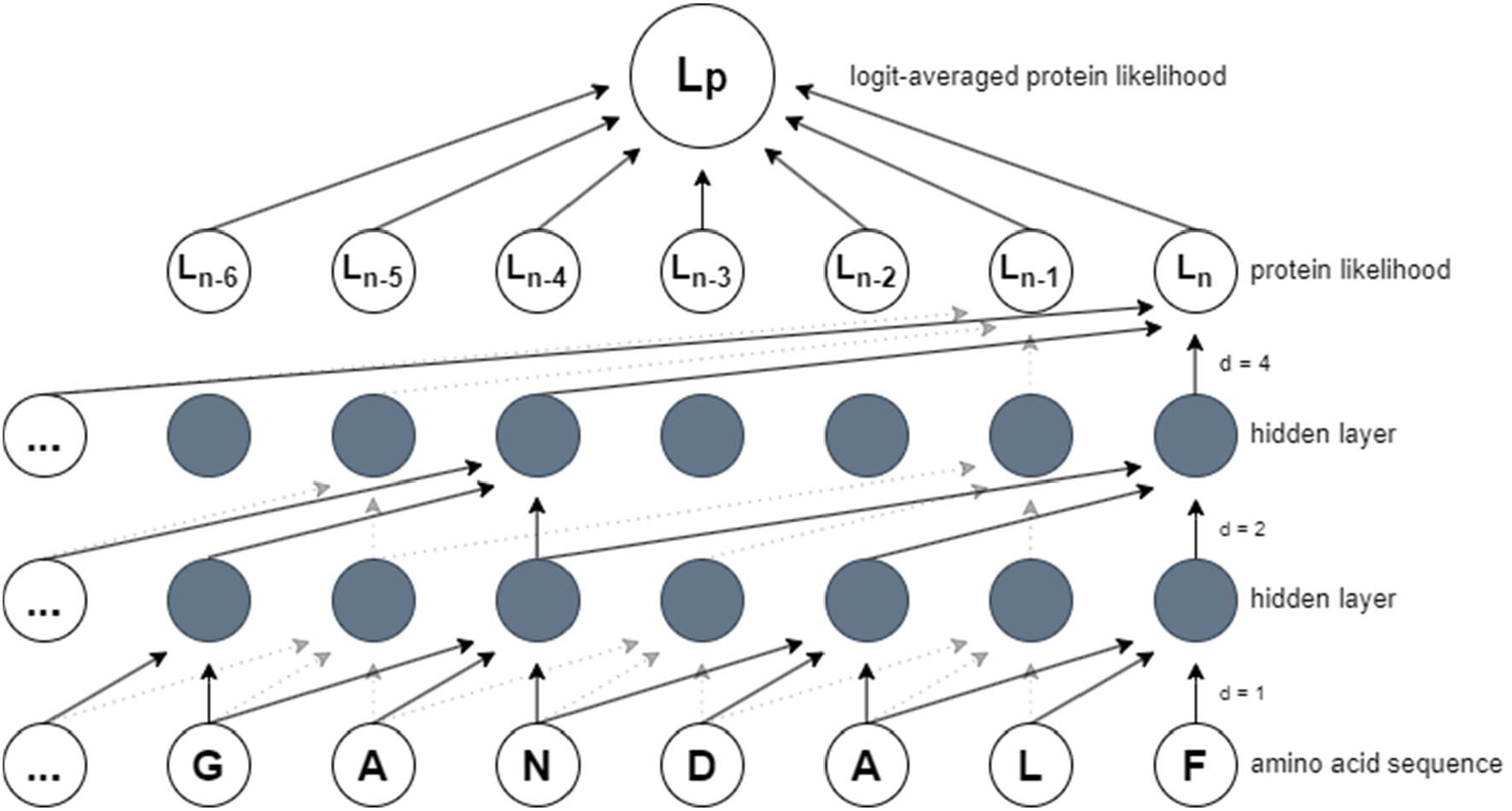
Model architecture. The pre-trained PSAURON TCN model produces a sequence of likelihoods for any length amino acid sequence. These likelihoods are logit-averaged into a single protein likelihood, L_p_, shown at the top. The simplified model shown here for demonstration purposes has 2 hidden layers, a kernel size of 3, and dilation factors of 1, 2, and 4. Left zero padding ensures that all layers are of equal size. The actual PSAURON TCN model uses 6 layers, 32 hidden units per layer, a kernel size of 16, and a dilation expansion factor of 2, for a total of 383,457 trainable parameters.

The PSAURON TCN belongs to a family of convolutional network architectures ^8^ that use dilated convolutions and left-zero padding to capture information from a wide receptive field, mapping input sequence data to an output of the same length. For our data type, amino acid sequences, the TCN architecture enables us to infer likelihood for any length protein. The PSAURON TCN (Figure 1) uses 6 layers, 32 hidden units per layer, a kernel size of 16, and a dilation expansion factor of 2, for a total of 383,457 trainable parameters. These hyperparameters were chosen to be similar to those used in Balrog, where Bayesian hyperparameter optimization was performed ^7,9^. To compare scores between proteins of varying lengths, we use the logit-averaged likelihood, a single number between 0 and 1 that represents an estimate of the predicted protein’s likelihood. Whereas the arithmetic average treats all likelihoods close to 0 or 1 as nearly identical, logit averaging more heavily weights likelihoods that are close to 0 or 1. For example, the arithmetic average of 0.500 and 0.999 is approximately 0.75, whereas the logit-average is approximately 0.97. Thus, if any part of a sequence appears protein-like, then PSAURON will be more likely to assign a high score. The logit-average for a sequence ***p*** of length ***n*** is defined as:

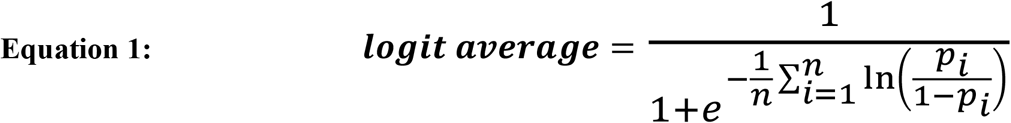

### Model Training

The PSAURON TCN model was trained on coding sequences (CDSs) from all plant and animal genomes available from the NCBI RefSeq database as of August 2023. These 1090 eukaryotic genomes included 171 plants, 359 invertebrates, 214 mammalian vertebrates, and 346 other vertebrates. Accession numbers for all genomes can be found in Supplemental Table 1. Sequences were excluded from analysis if they contained annotated in-frame stop codons, started with a non-standard start codon, contained non-ACGT nucleotides, or had a length that was not divisible by three. This selection process generated 33,869,774 unique CDSs.

Subsequences of 100 amino acids (aa) overlapping by 50aa (“shingles”) were extracted from protein sequences translated from annotated-frame and out-of-frame CDSs to generate positive and negative data respectively. For all shingles, start and stop codons were removed to prevent the learning algorithm from learning to label a sequence based on the presence or absence of those codons. To remain within memory limits during training, shingles were generated from 1,000,000 CDSs selected at random from the full CDS data set. From this subset, annotated-frame shingles numbered 11,406,219. The same number of out-of-frame shingles were randomly selected to maintain equal class balance during model training. The model was trained with binary cross-entropy loss with annotated-frame CDSs as positive data and out-of-frame CDSs as negative data, with a batch size of 1000. Adaptive moment estimation with decoupled weight decay regularization (AdamW) ^10^ was used to minimize loss during training. Model training required over 8 hours of computing time on a Google Colab server with an NVIDIA V100 16GB GPU.

### Protein Sequence Assessment

PSAURON assesses proteins by calculating a score between 0 and 1 for any given amino acid sequence, where 1 indicates that the sequence is a bona fide protein. The model is trained to maximize the difference in scores between sequences translated from in-frame CDS, which should receive scores close to 1, versus out-of-frame CDS, which should receive scores close to 0.Amino acid sequences that bear no resemblance to either in-frame or out-of-frame protein translations may receive unpredictable scores.

By default, PSAURON uses score from all six frames of the CDS to determine whether a sequence encodes a protein, and it intentionally ignores stop codons. Thus, a high PSAURON score does not guarantee that a sequence contains a valid ORF. Rather, because the model was trained to recognize both in-frame and out-of-frame translated sequences, scores from all frames are used to maximize predictive performance. For a sequence to be assessed as “positive,” the in-frame score must be greater than 0.5 and the mean out-of-frame score must be less than 0.5. These parameters may be changed by the user, and scores for all frames of all sequences are provided in the default output file. PSAURON can also run directly on protein sequence with no CDS provided, and in this case will provide only the score of the annotated frame.

### Proteome-wide Annotation Assessment

In addition to scoring each protein individually, PSAURON also provides an overall score between 0 and 100 for the entire input annotation. This score represents the percentage of proteins provided in the annotation that receive a “positive” assessment as described above. The whole-proteome PSAURON score can change depending on user-defined parameters that trade sensitivity for precision. The whole-proteome score as well as scores for each annotated protein sequence are provided in the default output file, and these may be used to sort proteins by PSAURON’s confidence.

## RESULTS

Table 1 provides whole-proteome PSAURON scores for a variety of model organisms including animals, plants, fungi, and prokaryotes. Supplemental Tables S2-S3 contain PSAURON scores for all proteins in the Matched Annotation from NCBI and EMBL-EBI (MANE) annotation as well as the UniProt reference rice proteome. For these evaluations PSAURON was run in single-frame or protein mode with a score cutoff of 0.5 and no minimum protein length threshold.

**Table 1.**
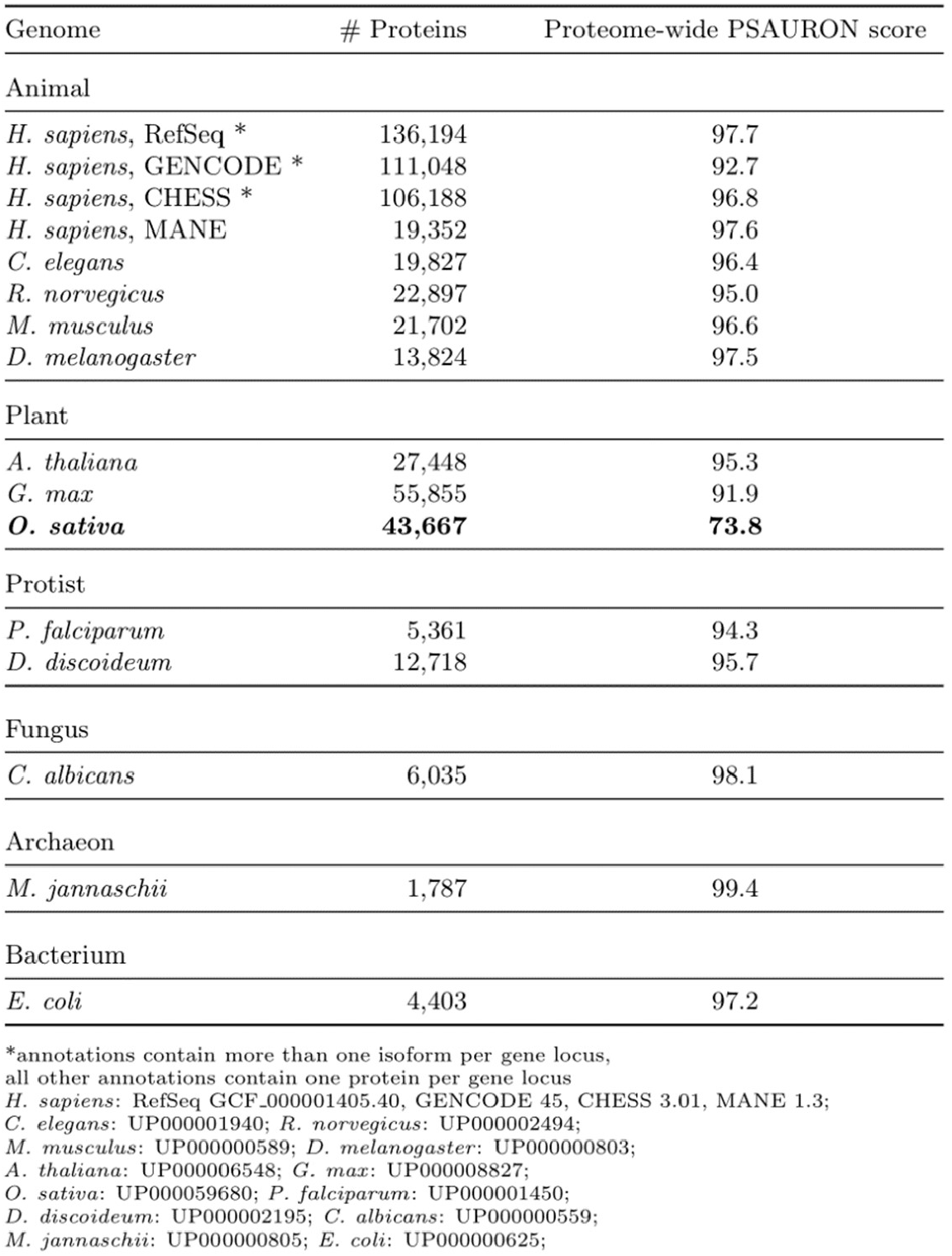
Reference scores. Proteome-wide PSAURON scores for selected UniProt reference proteomes and human genome annotation databases. PSAURON was run in protein mode with the default score threshold of 0.5; the proteome-wide PSAURON score in this table represents the percentage of proteins that scored above 0.5. All proteome-wide PSAURON scores arc greater than 90 with the sole exception of rice, *O. sativa*, in bold.

Interestingly, despite PSAURON having been trained exclusively on eukaryotic proteins, the highest score, 99.4%, was achieved by a low-GC-content archaeon, *M. jannaschii*. Prokaryotic annotation in low-GC genomes is highly accurate, and PSAURON agrees with the annotation for nearly all the proteins in this genome. All other reference proteomes achieved scores greater than 90%, with the sole exception of *Oryza sativa subsp. japonica* (rice), which scored 73.8%. This whole-proteome score suggests that nearly 30% of the annotated proteins in rice might be incorrect.

### Comparison of PSAURON scores and protein structure predictions

The AlphaFold2 program has been demonstrated to be remarkably accurate at predicting the three-dimensional structure of many proteins ^11,12^. AlphaFold2 also produces a score for each structure, known as the predicted Local Distance Difference Test (pLDDT), which indicates the program’s confidence in its structural prediction. High-scoring structures are likely to be correct, while lower-scoring structures, with scores below 70, are “considered low confidence and should be treated with caution.” ^11^ Not all proteins fold into a stable structure or receive a high pLDDT from AlphaFold2; e.g., for the human genome about 58% of proteins fall into the high-confidence group, with scores above 70 ^13^. Despite these caveats, the accuracy of AlphaFold2 makes it a powerful tool for independently evaluating the quality of a set of predicted proteins, as we have demonstrated previously ^14^.

For the experiments here, we hypothesized that the scores assigned by AlphaFold2 should follow a similar distribution across a wide variety of organisms. Figure 2 shows the distributions of pLDDT scores for five eukaryote genomes, including two animals (human and mouse), two plants (rice and the model plant *Arabidopsis thaliana*), and one fungus. As expected, all but one of the distributions look similar. The sole outlier in this analysis is the rice proteome, shown in red, which has a secondary peak in pLDDT density near 60, while all other proteomes peak near 90.

**Figure 2:**
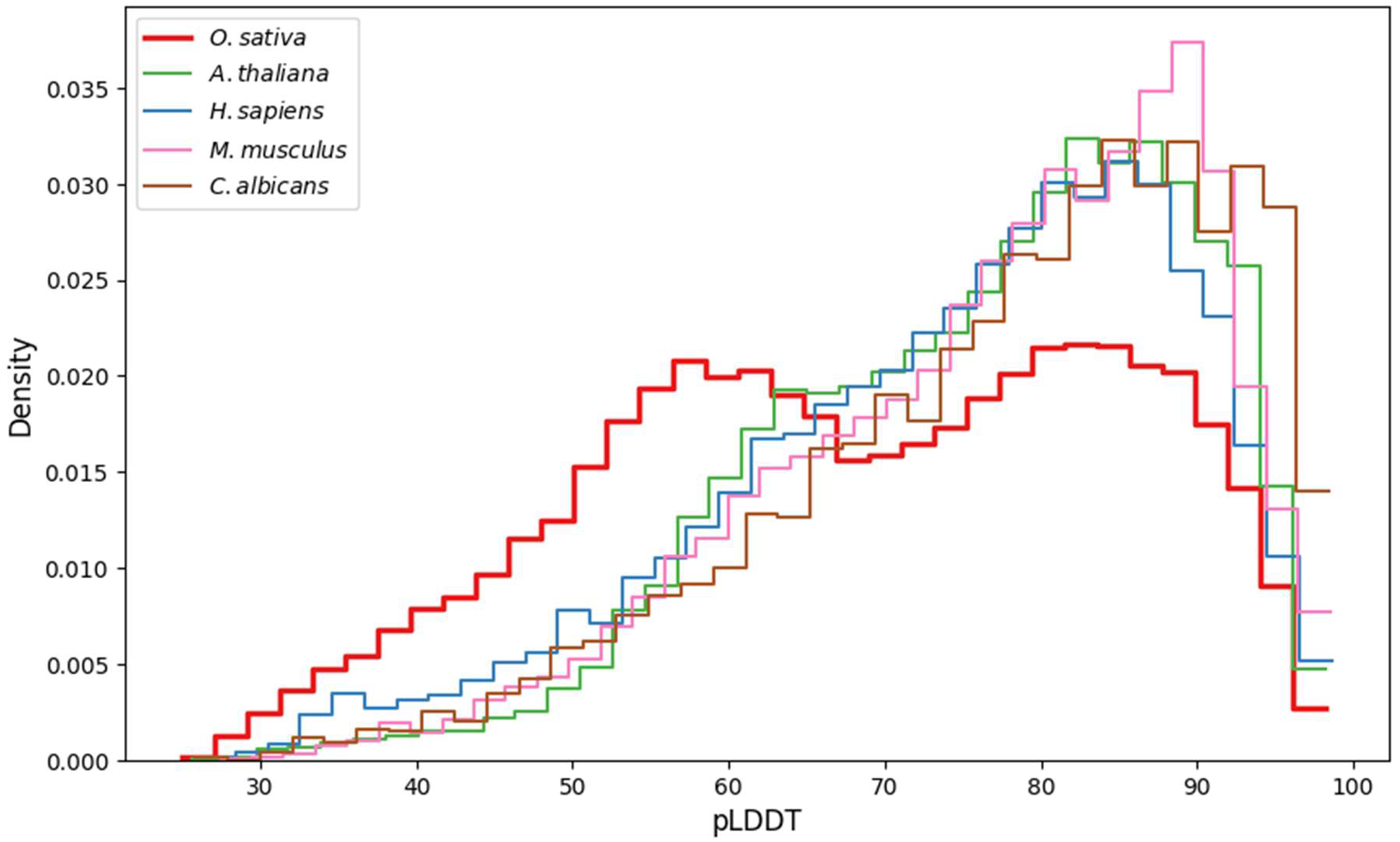
AlphaFold2 confidence distribution of reference proteomes. Histograms of AlphaFold2 pLDDT score of proteins in UniProt reference proteomes. High-quality eukaryotic proteomes tend to show similar distributions of scores. The rice proteome is an outlier in pLDDT distribution.

Worth noting is that we obtained this low proteome-wide PSAURON score for rice on the UniProt reference proteome annotation, which is based on the Rice Annotation Project Database ^15,16^, but not for the RefSeq rice annotation, which we scored at 96.0. Both annotations use the same rice genome assembly, which suggests the issue with the rice proteome is a result of flawed annotation rather than assembly.

As shown in Figure 3, rice proteins with low PSAURON scores also tended to receive low AlphaFold2 confidence scores, with 96.9% of PSAURON’s low-scoring proteins receiving a pLDDT below 70. Note that PSAURON was trained in an unsupervised manner on CDS sequences alone and was given no information on protein structure. Thus, in combination with unusually low structure scores, the current UniProt rice proteome annotation appears to contain several thousand proteins that may be mis-annotated. Interestingly, if one were retain only the proteins to which PSAURON assigned a high score, then the distribution of AlphaFold2 scores for rice (blue curve in Figure 3) would more closely resemble the other distributions shown in Figure 2.

**Figure 3:**
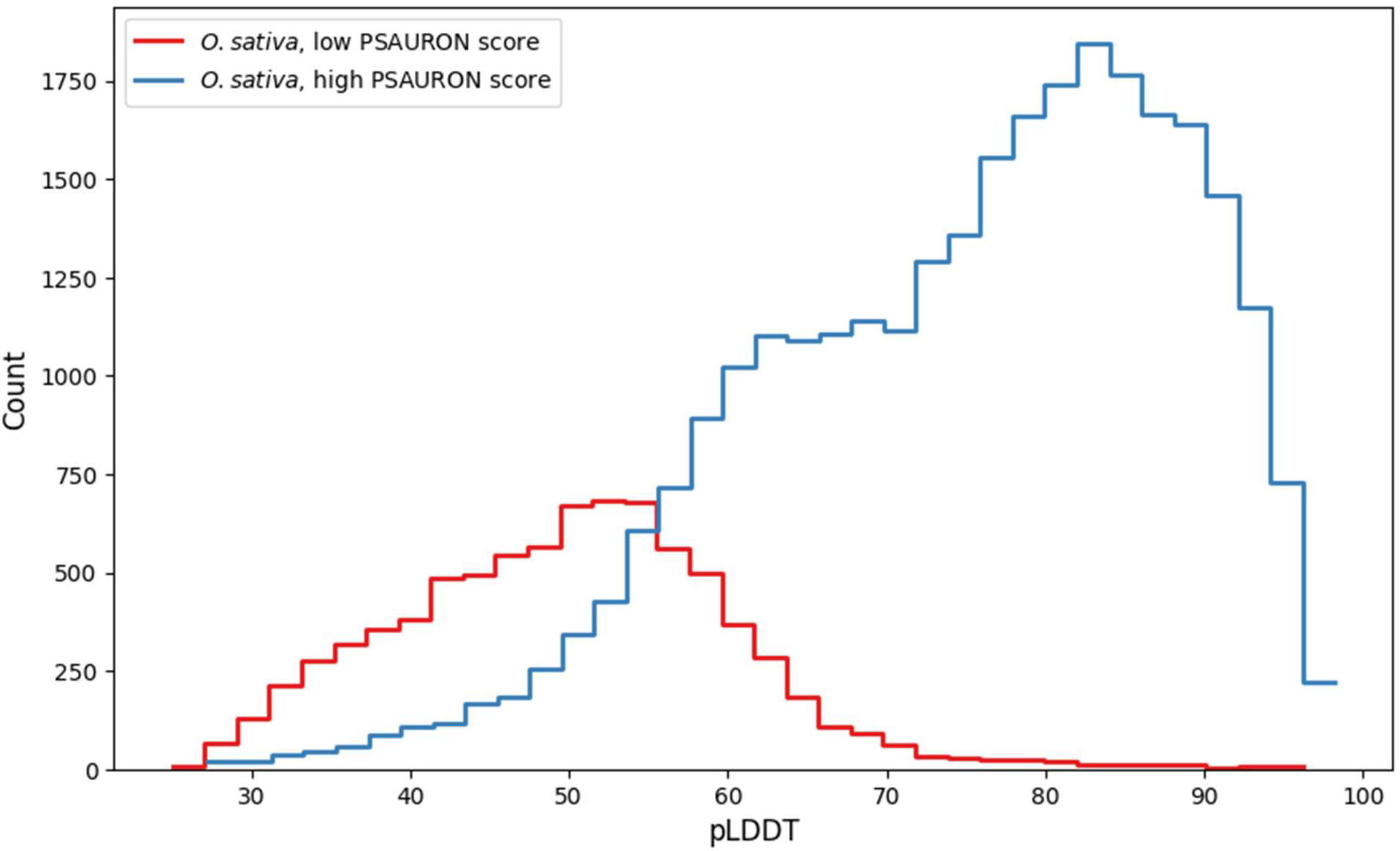
Rice reference proteome AlphaFold2 predictability separated by PSAURON score. Histograms of AlphaFold2 pLDDT scores of the rice reference proteome. The blue curve shows proteins with high PSAURON scores (> 0.8) and the red curve shows proteins with low PSAURON scores (< 0.2). 96.9% of all low-PSAURON-scoring proteins receive a pLDDT below 70. The pLDDT distribution of high-PSAURON-scoring proteins in the rice proteome more closely resembles those of other high-quality reference proteomes.

### Consistency with manual curation (Arabidopsis)

The Arabidopsis Information Resource (TAIR) provides a curated ranking system for genes in the *A. thaliana* reference genome, ranking them from zero to five stars based on transcriptomic and proteomic data as well as multiple sequence alignment and genomic conservation analysis ^17^. Using the TAIR10 data, we scored all proteins with PSAURON in single-frame mode to generate PSAURON score distributions for each ranking group. Using a consumer-grade Dell XPS 13 laptop running Ubuntu 20.04 LTS with a 4-core Intel i7-1185G7 at 3.00 GHz, PSAURON scored the full TAIR10 set of 35,386 proteins in 132 seconds with a peak memory usage of 1.95 GB.

The PSAURON scores were consistent with the TAIR confidence rankings: as shown in Figure 4, the higher-confidence proteins (2-5 stars) received very high PSAURON scores with a tight distribution of values close to 1.0. For low-confidence proteins with one- or zero-star confidence ratings, PSAURON produced lower average scores with a wider distribution, particularly for the zero-star proteins. Thus, the independently-determined manual curation of *Arabidopsis* supports the scores assigned by PSAURON.

**Figure 4:**
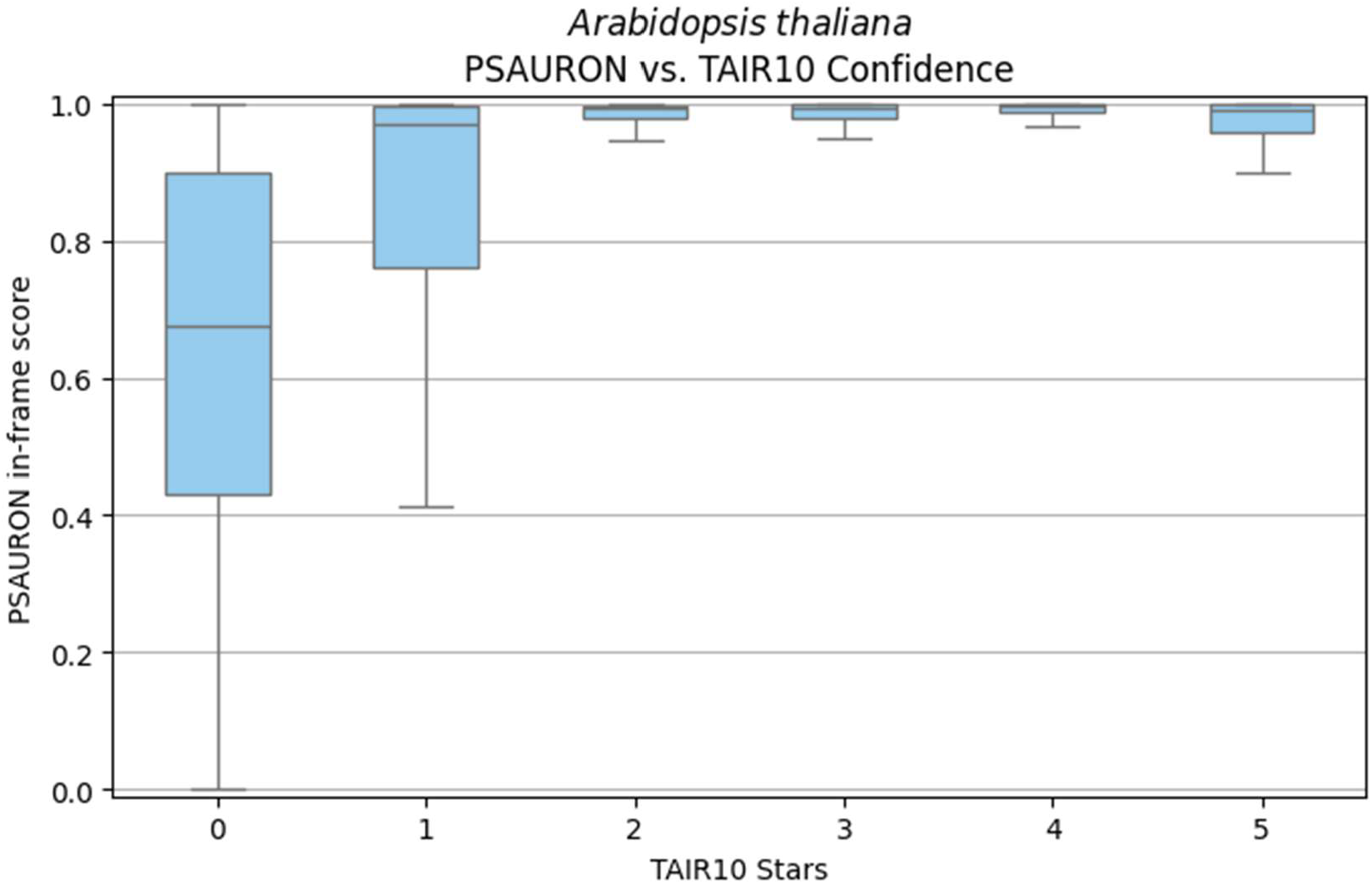
PSAURON scores for Arabidopsis proteins by TAIR10 confidence. Box and whisker diagram of the PSAURON score distribution for annotated coding sequences in the TAIR10 reference genome. The x-axis shows the TAIR confidence ranking for each group of genes, ranging from a minimum of zero (low confidence) to a maximum of 5 stars. The boxes show the interquartile range (IQR) of individual protein PSAURON scores. The whiskers extend to the furthest point within 1.5 times the IQR.

## DISCUSSION

PSAURON is a computational tool that can recognize protein-coding genes in a wide range of eukaryotic species. Similar to Balrog ^7^, and in contrast to most previous *ab initio* gene finders, PSAURON recognizes features of proteins rather than DNA sequences. Thus, unlike most gene finders, PSAURON does not need to be trained specifically on each species, but only needs to be trained once, as we have done here using a diverse dataset derived from over 1000 plant and animal genomes. PSAURON can evaluate protein sequences and provide confidence scores that reflect the predicted likelihood of each annotated protein with no need for organism-specific retraining.

PSAURON can be used to score all proteins in a genome individually and to produce a single overall score. This represents a novel approach for quality assessment of genome annotation in eukaryotes, one that does not require the prior identification of a set of single-copy genes or other species-specific input. This approach addresses a critical need in the field of genomics, as the accuracy of protein-coding gene annotations is crucial for downstream analyses and applications. As demonstrated by the thousands of low-quality proteins present in the reference rice proteome, problems with automated annotation workflows may go unnoticed even in the most important model species.

PSAURON can also generate model-based confidence scores for individual proteins, requiring no additional experiments such as proteomics or large-scale manual curation. Curated confidence rankings, such as those available for Arabidopsis, are not feasible to produce for every genome sequenced today. The relationship between PSAURON scores and the TAIR10 gene confidence rankings for Arabidopsis suggests that PSAURON scores reflect protein annotation quality. By providing a confidence score for each annotated gene, PSAURON can enable researchers to quickly identify potential inaccuracies and prioritize sequences that require either further validation or removal from annotation. PSAURON is not intended to be the sole oracle of annotation quality. Instead, we hope PSAURON will provide additional useful information to researchers wishing to produce accurate annotations for a wide range of species.

One unexpected result of our evaluations was that the annotation of rice, arguably humanity’s most important food source ^18^, received a substantially lower proteome-wide PSAURON score compared to other reference proteomes. This suggests that the current reference rice proteome contains thousands of inaccurate protein annotations. This finding was further supported by the observation that low-PSAURON-scoring rice proteins tend to have low-confidence AlphaFold2 structural predictions, as represented by the abnormal pLDDT distribution of the rice proteome. We have previously shown that pLDDT can be used as a metric to help improve protein annotations, even in highly-studied genomes ^14^. It is important to note that low pLDDT scores alone are not definitive proof that sequences do not represent real proteins, particularly for those proteins that are inherently unstructured. Still, given the substantially skewed pLDDT distribution of low-PSAURON-scoring proteins in the rice proteome, our results suggest the rice annotation contains an unusually high number of errors.

Even in reference annotations for important model species, technical and biological noise remains a problem ^19^. The Matched Annotation from NCBI and EMBL-EBI (MANE) is a high-quality human annotation that has been proposed as the “universal standard” for clinical variant reporting ^20^. As expected, MANE received one of the highest scores among all annotations tested here, with 97.6% of included proteins receiving a positive PSAURON assessment. In some cases, users may wish to include more proteins in an annotation at the cost of including technical and biological noise ^19^. The GENCODE human annotation ^21^, for example, aims to be a comprehensive view of human biology, while MANE restricts itself to a single high-confidence transcript per protein-coding locus ^20^. Accordingly, MANE receives a higher PSAURON score than GENCODE. A higher score does not necessarily mean that one database is superior, as it may be the result of logical decisions based on different use cases for each database. Even so, some MANE proteins may need correction ^14^. Only 83% of 234 “uncharacterized” proteins (i.e., proteins with no known function) in MANE v1.3 scored well according to PSAURON. Additionally, 2 out of 3 “umcharacterized” proteins in MANE v1.3 scored poorly according to PSAURON, suggesting that even the most highly curated human gene set is not yet perfect.

Our experiments demonstrated the effectiveness of PSAURON through a series of analyses on reference datasets. As expected, the proteome-wide PSAURON scores for a variety of highly-studied model organisms were high, commensurate with the expected quality of their annotations. Perhaps surprisingly, PSAURON also appeared to be effective for scoring annotation in the bacterium *Escherichia coli* and the archaeon *Methanocaldococcus jannaschii*, although its training data did not include organisms from either kingdom. In contrast to eukaryotic genomes, prokaryotic genomes are relatively simple to annotate, particularly when they have low GC-content ^7^. These high scores suggest that PSAURON, which was trained exclusively on eukaryotic protein sequence, learned to recognize more universal features of protein sequences, features that apply to all living species.

In its current form, PSAURON is not intended to be used as an *ab initio* gene finder, because it does not consider all possible combinations of exons and introns in a genomic DNA sequence as some *ab initio* gene finders do ^22–25^. Because it scores a user-provided annotation, PSAURON cannot estimate the completeness of that annotation. Rather, it can be used as a quality-control measure, serving as an estimate of annotation noise and a means to highlight potentially inaccurate annotations. However, PSAURON could be used as a module in a more general *ab initio* gene finder in future work.

In conclusion, PSAURON represents a new method for assessing the accuracy of protein-coding gene annotation across a wide range of eukaryotic species. Its ability to leverage a large, diverse training dataset to assess the quality of protein-coding gene annotations has the potential to improve the accuracy and reliability of protein-coding gene annotation.

## ACKNOWLEDGEMENTS

The authors would like to thank all two and four legged members of the Salzberg and Pertea labs, as well as members of the Battle lab including Josh Weinstock for assistance with statistical thinking. This work was supported in part by NIH grants R01-HG006677 and R35-GM130151.

## Contributions

MJS designed the architecture of PSAURON and wrote the code. MJS and SLS were responsible for initial project conceptualization and general design details. MJS, AVZ, and SLS designed the computational experiments. MJS ran all experiments. MJS, AVZ, and SLS wrote and edited the manuscript.

